# MeDEStrand: an improved method to infer genome-wide absolute methylation level from DNA enrichment experiment

**DOI:** 10.1101/194431

**Authors:** Jingting Xu, Shimeng Liu, Ping Yin, Serdar Bulun, Yang Dai

## Abstract

**Background:** DNA methylation of dinucleotide CpG is an essential epigenetic modification that plays a key role in transcription. Bisulfite conversion method is a “gold standard” for DNA methylation profiling that provides single nucleotide resolution. However, whole-genome bisulfite conversion is very expensive. Alternatively, DNA enrichment-based methods offer high coverage of methylated CpG dinucleotides with the lowest cost per CpG covered genome-wide and have been used widely. They measure the DNA enrichment of methyl-CpG binding, therefore do not directly provide absolute methylation levels. Further, the enrichment is influenced by confounding factors besides the methylation status, e.g., CpG density. Computational models that can accurately derive the absolute methylation levels from the enrichment data are necessary.

**Results:** We present ‘MeDEStrand’, a method uses sigmoid function to estimate and correct the CpG bias from the numbers of reads that fell within bins that divide the genome. In addition, unlike the previous methods, which estimate CpG bias based on reads mapped at the same genomic loci, ‘MeDEStrand’ processes the reads for the positive and negative DNA strands separately. We compare the performance of ‘MeDEStrand’ with three other state-of-the-art methods ‘MEDIPS’, ‘BayMeth’ and ‘QSEA’ on four independent datasets generated using immortalized cell lines (GM12878 and K562) and human patient primary cells (foreskin fibroblast and mammary epithelial). Based on the comparison between the inferred absolute methylation levels from MeDIP-seq and the corresponding RRBS data, ‘MeDEStrand’ shows the best performance at high resolution of 25, 50 and 100 base pairs.

**Conclusions:** ‘MeDEStrand’ benefits from the estimation of CpG bias with a sigmoid function and the procedure to process reads mapped to the positive and negative DNA strands separately. ‘MeDEStrand’ is a tool to infer whole-genome absolute DNA methylation level at the cost of enrichment-based methods with adequate accuracy and resolution. R package ‘MeDEStrand’ and its tutorial is freely available for download at https://github.com/jxu1234/MeDEStrand.git

## Background

DNA methylation of dinucleotide CpG is an essential epigenetic modification that plays a key role in transcription regulation. Sequencing-based DNA methylation profiling techniques include whole-genome bisulfite sequencing (WGBS), reduced-representation bisulfite sequencing (RRBS) and enrichment-based methods, such as methylated DNA immunoprecipitation followed by sequencing (MeDIP-seq) and methyl-CpG binding domain protein sequencing (MethylCap-seq/MBD-seq), etc. These methods generate informative and comparable results, yet differ significantly in the extent of genomic CpG coverage, resolution, quantitative accuracy and cost [1, 2].

WGBS and RRBS are the “gold standard” methods for DNA methylation studies [3, 4]. They yield single-nucleotide resolution of the methylation status in the scale from 0 to 1. RRBS provides substantial coverage of CpGs in CpG islands, with lower CpG coverage genome-wide, while WGBS offers greater CpG coverage genome-wide but significantly higher cost.

On the other hand, methylated DNA enrichment-based methods, such as MeDIP-seq and MethylCap-seq/MBD-seq, offer high coverage of methylated CpG dinucleotides and have the lowest cost per CpG covered genome-wide [1]. They measure the enrichment of methylated DNA fragments that can be used to infer regional methylation status. However, the absolute methylation levels must be derived from proper computational models that eliminate the effects of confounding factors. It has been shown that MeDIP-derived data need to be corrected for CpG density effects to obtain the unbiased methylation levels [5, 6].

Several methods have been developed to infer absolute DNA methylation levels from the enrichment-based DNA methylation profiles. ‘MEDME’ is one of the earlier methods to estimate DNA methylation levels based on the microarray derived MeDIP-enrichment (MeDIP-chip) experiment on normal human melanocytes [6]. Treated with CpG methyltransferase (M.SssI, NEB), the genomic DNA fragments were fully methylated. ‘MEDME’ assumed that only methylated CpGs generated the enrichment of antibody-binding fragments. Based on the observed sigmoidal relationship between the enrichment signal and methylated CpG density, a four-parameter logistic model was used to fit the observed data and establish the relationship between the enrichment signal and methylated CpG density (Figure 1(A)). The estimated model is assumed to be generic and can be applied on other cells to infer absolute DNA methylation levels. The ‘MEDME’ model was developed based on microarray derived MeDIP-enrichment data and has not been extended to sequencing-based MeDIP-seq data. The inferred absolute methylation level for a 1 kb window is at a low resolution.

**Figure 1.**
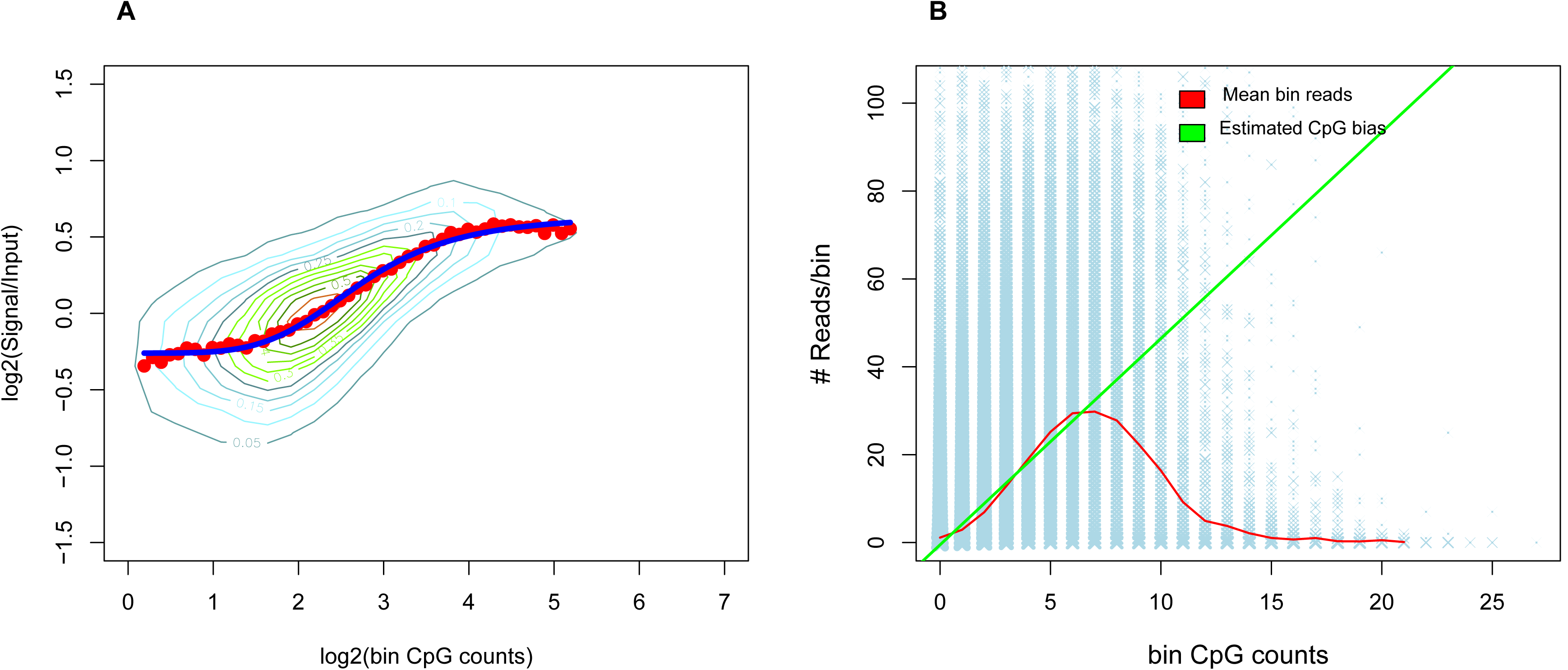
‘MEDME’ experiment versus ‘MEDIPS’ calibration plot. **A** The sigmoidal relationship between microarray-based MeDIP-chip signal and number of fully methylated CpGs was revealed by ‘MEDME’ experiment in log-scale. The red dots identify the median MeDIP logR within each window across the dynamic range of numbers of methylated CpGs. **B** ‘Calibration’ plot used in ‘MEDIPS’ method to estimate CpG bias from the bin reads. The blue strips show the plots of the bin reads vs. bin CpG counts. The relationship between the means of the bin reads and bin CpG counts are shown as the red line. The green line comes from fitting a simple linear regression of the mean bin reads at low CpG density regions.

‘BATMAN’ is another method for inferring absolute methylation level for array-based MeDIP-chip data [5]. It also provides a modified model that theoretically extends its application to sequencing-based MeDIP-seq data. ‘BATMAN’ assumes a linear relationship between MeDIP-chip signal and CpG density and models the observed signals as the additive results of ‘base effects’ and CpG methylation levels. ‘BATMAN’ uses Bayesian inference to estimate the posterior distribution of the methylation parameters given the MeDIP-chip signals, thus generates the absolute methylation level profile. It is recommended in ‘BATMAN’ to use 100 bp bin for the inference of absolute methylation levels. Due to its relatively complex Bayesian inference based on ‘nested’ sampling algorithm, ‘BATMAN’ was reported to be time consuming. Later, ‘MEDIPS’ was developed for the sequencing-based enrichment MeDIP-seq experiment [7, 8]. ‘MEDIPS’ is a time-efficient statistical method that combines the findings from ‘MEDME’ and ‘BATMAN’. It adapts the concept proposed in ‘BATMAN’ that the main confounding factor comes from CpG density. To estimate the CpG density effects (i.e., bias), ‘MEDIPS’ divides genome into 50 or 100 bp bins and counts the reads fell in the bins (a.k.a. bin reads), groups bins according to their CpG counts, then uses simple linear regression to model the relationship between the bin reads and CpG counts of the bins. The fitted line is used as an estimate of the CpG density bias, which is subsequently corrected from the observed bin reads. The corrected bin reads scaled to values between 0 to 1 are reported as the inferred genome-wide bin-based absolute methylation levels.

Recently, new methods were developed to incorporate additional experimental data to improve the accuracy for the inference of the absolute methylation levels. ‘BayMeth’ is an empirical Bayes approach that uses a fully methylated control sample to transform observed bin reads into absolute methylation levels [9]. ‘methylCRF’, a novel Conditional Random Fields-based algorithm that integrates methylated DNA immunoprecipitation (MeDIP-seq) and methylation-sensitive restriction enzyme (MRE-seq) sequencing data to predict DNA absolute methylation levels at single-CpG resolution [10]. Lately, a new method ‘QSEA’ was developed to improve ‘BayMeth’ [11]. ‘QSEA’ constructs two virtual control samples, i.e., ‘TCGA’ and ‘bind’, to provide good performance without the use of experimental control samples.

Despite the improvement acquired by ‘MEDIPS’ following the development of ‘MEDME’ and ‘BATMAN’, there is still room for further improvement. Firstly, ‘MEDIPS’ uses a simple linear regression model to estimate the CpG density bias. It is observed that at low CpG density regions, the means of bin reads increase when CpG counts of the bins increase. Based on the information that CpGs are mostly methylated at low CpG density regions, ‘MEDIPS’ thus, considers that it is the CpG counts rather than the methylation levels (which are viewed as fixed) of the bins that contribute to the increment of the bin reads. On the other hand, the bin reads began to decrease after reaching the summit and continued the decrease extending to the high CpG density regions. High CpG density regions correspond to CpG Islands or high CpG promoter regions that are mostly hypo- or un-methylated. So, the decreasing trend of the bin reads is the results of significant effects of decreased methylation level (although the CpG density remains high). To estimate the CpG bias for all regions, ‘MEDIPS’ uses a simple linear regression model to fit the means of bin reads at low CpG regions and extrapolates the fitted line to the high CpG regions assuming the CpG effects be non-decreasing (Figure 1(B), green line). The fitted line, an estimate of CpG bias, is used by ‘MEDIPS’ to correct CpG bias from the bin reads.

The ‘MEDME’ experiment reveals the true relationship between the CpG density and its influence on the MeDIP signals. ‘MEDME’ constructed a CpG methyltransferase (M.SssI, NEB) treated sample of which all CpGs are full methylated. Since CpG methylation is controlled at the same, the experiment reveals that the signal dependency on the CpG density is not linear but sigmoidal (Figure 1(A)). Therefore, by the extrapolation of the fitted line at low CpG density regions to the high CpG density regions, ‘MEDIPS’ does not take into consideration the saturation effect of methyl-CpG binding, which may lead to over-estimation and over-correction of the CpG bias at high CpG density regions. Note, although the ‘MEDME’ experiment was based on microarray-derived signals (e.g., the signals and bin CpG counts were log-transformed), it can be revealed that the inverse log transformation of these values maintains a sigmoidal relationship except being a little flatter. Thus, we postulate that the CpG density effect is sigmoidal in the MeDIP-seq context.

Secondly, none of the current methods paid attention to the effects of asymmetric CpG methylation, i.e., methylation of cytosine in the ‘CG’ context on one DNA strand and un-methylated adjacent cytosine in the ‘GC’ context (or still ‘CG’ from 5 end to 3 end) on the other DNA strand (Figure 2). Our investigation of the RRBS data of GM12878 cell line shows that cytosine methylation in the dinucleotide CpG on the positive DNA strand and negative DNA strand is slightly different (Figure 3). We found that adjacent cytosine methylation in the CpG pair on the positive and negative DNA strands are highly concordant genome-wide with Pearson correlation coefficient (PCC) being ∼ 0.97. However, when genome is divided into bins of various sizes that contain consecutive CpGs (Figure 2), we observed that bin-based DNA methylation levels (the mean of methylation levels of all the CpGs in the bin) for the positive and negative DNA strands have decreased concordance when the bin size increases (Table 1). Six chromosomes (1, 2, 11, 12, 21, 22) were selected to represent chromosomes of large, medium and small sizes. The complete results are provided in the supplementary **Table S1**.

**Figure 2.**
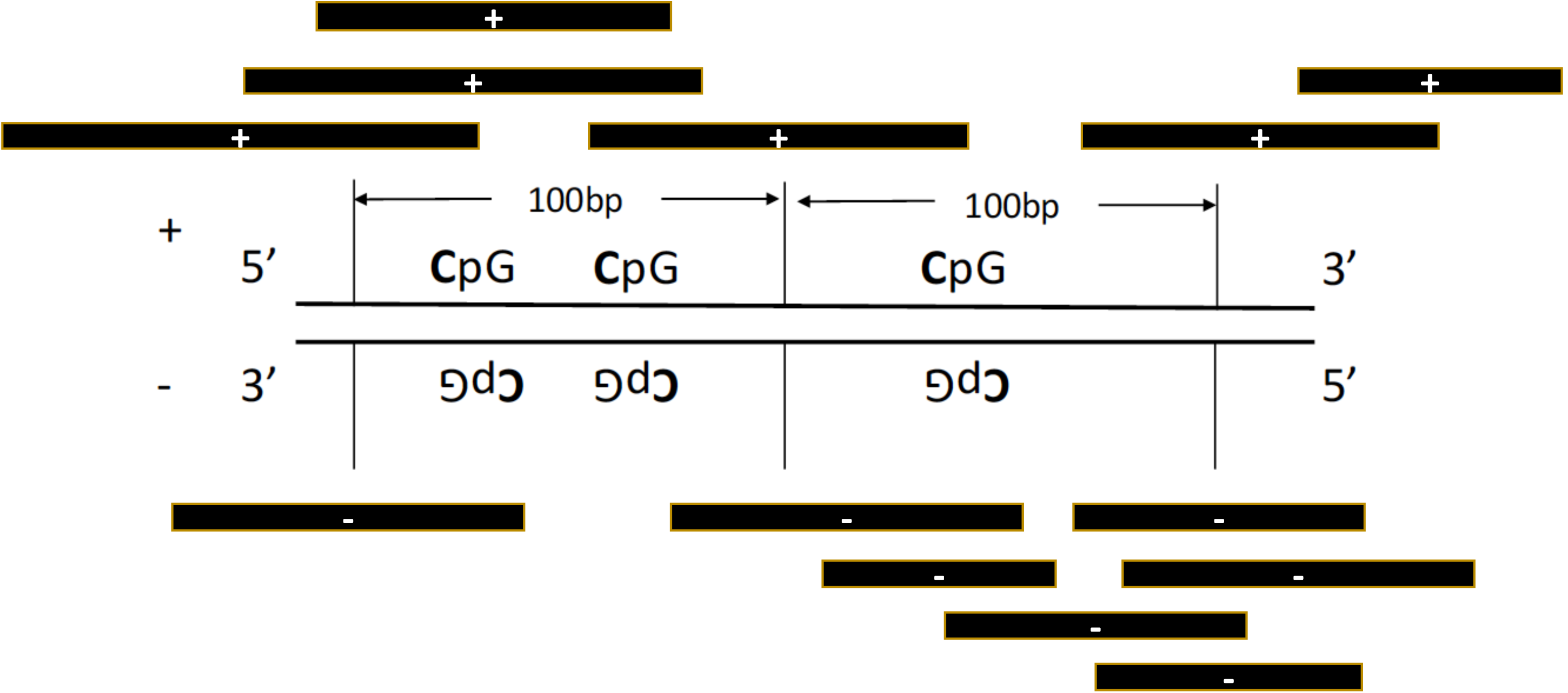
Illustration of 100bp bins on a segment of genome and the numbers of reads fell in the bins (bin reads). Bin reads measures the Methyl-CpG binding signal strength at the loci. Reads from both DNA strands are usually combined for bin reads for the loci. In our method, bin reads are counted for the positive and negative DNA strands separately.

**Figure 3.**
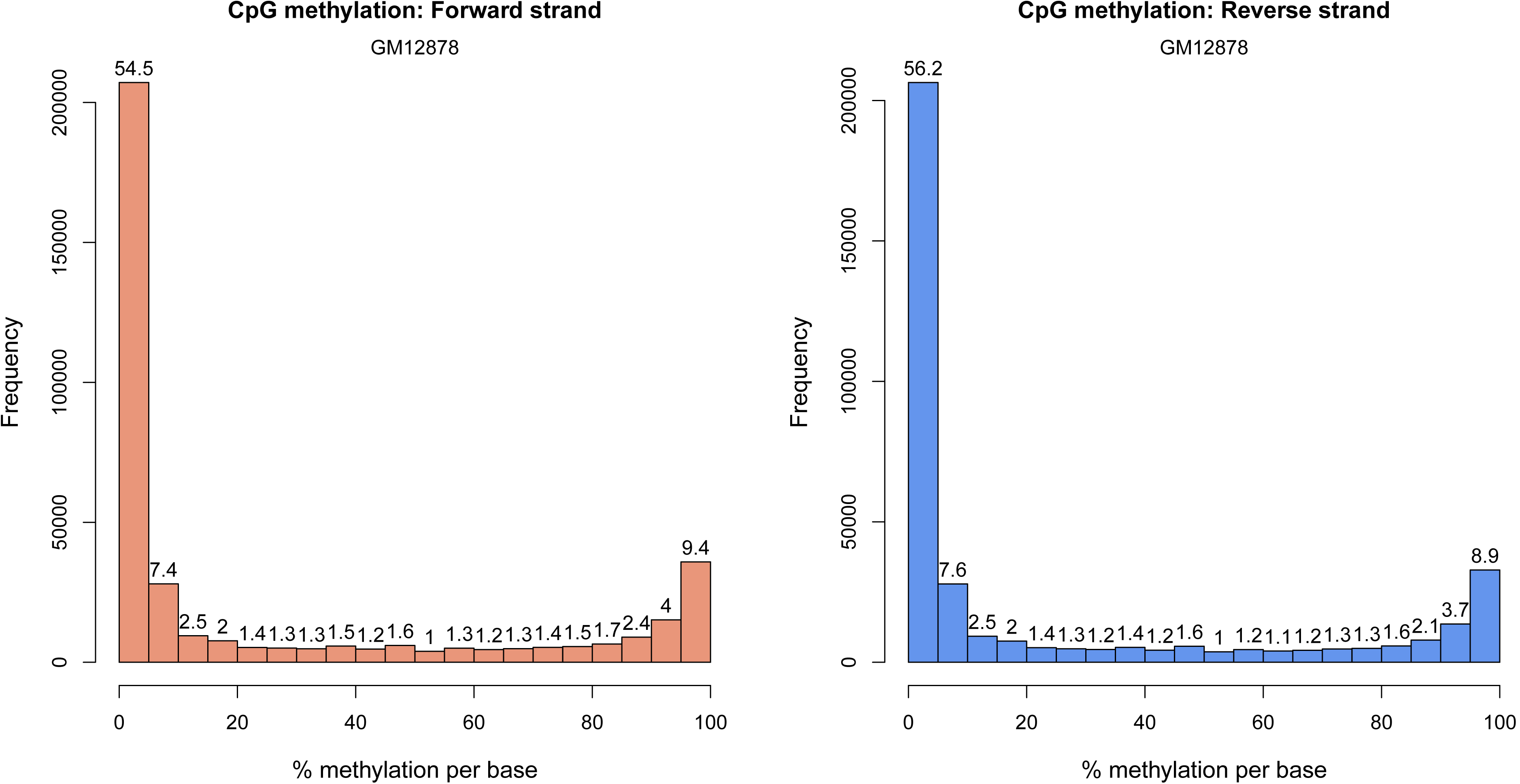
Histogram of cytosine methylation levels within CpGs on the forward (positive) and reverse (negative) DNA strands from the RRBS data of cell line GM12878.

**Table 1.** Pearson correlation coefficient of the bin methylation levels between the positive and negative DNA strands at various bin size (bp). Cell line: GM12878, Data: RRBS

| bin size | chr1 | chr2 | chr11 | chr12 | chr21 | chr22 |
| --- | --- | --- | --- | --- | --- | --- |
| 25 | 0.96 | 0.95 | 0.96 | 0.96 | 0.96 | 0.96 |
| 50 | 0.93 | 0.92 | 0.93 | 0.94 | 0.94 | 0.94 |
| 100 | 0.88 | 0.87 | 0.87 | 0.89 | 0.89 | 0.88 |
| 150 | 0.86 | 0.86 | 0.85 | 0.88 | 0.85 | 0.85 |
| 200 | 0.86 | 0.85 | 0.84 | 0.87 | 0.85 | 0.84 |

Since the average CpG methylation levels within the bins reflect regional methylation status, which results in the abundance of regional methyl-CpG binding, this discordance at bin level between the two DNA strands implies its possible effects on the MeDIP enrichment in a strand-specific fashion.

Based on the analysis described above, we propose our method ‘MeDEStrand’ (Inferring genome-wide absolute **me**thylation level from **D**NA **e**nrichment experiment utilizing **strand**-specific information) for the inference of absolute methylation level. ‘MeDEStrand’ improves ‘MEDIPS’ in the following two aspects:

- To use a logistic regression model for the estimation of the CpG bias. As shown in the ‘MEDME’ experiment, the upper asymptote of the logistic regression function is more suitable to model the saturation point of methyl-CpG-binding in high CpG density regions.
- To estimate and correct CpG bias for the positive and negative DNA strands separately to accommodate asymmetric/strand-specific methylated DNA enrichment.

## Methods

### Materials

MeDIP-seq and RRBS data used in our model development are from the ENCODE Consortium [12]. To test the genericity of our model, all cell types with both the MeDIP-seq and RRBS data available could be used. The RRBS data were used as a ‘gold standard’ for the method validation and comparison in the previous published methods.

However, to limit variation that could be introduced by the heterogeneity of cell types in tissue, only immortalized cell lines (GM12878 and K562) and primary cells (foreskin fibroblast and mammary epithelial) were selected in the study to provide high consistency between the RRBS and MeDIP-seq data. Since DNA methylation is a highly dynamic and transient epigenetic event [13-15], these datasets are most matched thus provide high confidence of the results and the conclusions.

The MeDIP-seq data in the ‘. bam’ format and the RRBS data in the ‘. sra’ format were downloaded from the ENCODE Consortium [12]. The ‘raw’ RRBS data (in the ‘. sra’ format) enables us to retrieve the methylation value for every single CpG cytosine such that the strand-specific CpG methylation information can be investigated. SRA Toolkit [16], samtools [17], Bismark [18] and Bioconductor packages methylKit [19], IRanges [20] were used in the data analysis.

### Model and the algorithm

Our method was motived by the ‘MEDME’ experiment. ‘MEDME’ constructed a fully methylated sample to establish the one-to-one relationship between MeDIP-chip signals and the corresponding numbers of methylated CpGs. MeDIP signal is explained by only one factor, i.e., the number of methylated CpG.

Since the cell was treated and all CpGs are methylated, we view the sigmoidal signal increase from the CpG density increase as the CpG bias since the increase only depends on CpG density change (Figure 1(A)). The CpG bias can be estimated and eliminated from the observed signals. For the same reason as ‘MEDIPS’ that in the high CpG density regions, CpGs are likely to be hypo- or un-methylated, CpG density does not act as the sole factor affecting the enrichment signal. We use a sigmoidal logistic regression model to fit the means of bin reads from the low CpG density regions, let its upper asymptote be the maximum mean observed and extend the estimate to the high CpG density regions as the CpG bias for all the regions (Figure 4, blue line). Based on the above reasoning, we propose to model the bin reads (y) to be the results of CpG methylation induced signal (*M*_*CpG*_) multiplied by the CpG density effect (*f*(*n*_*CpG*_)) which is a function of the bin CpG counts:

**Figure 4.**
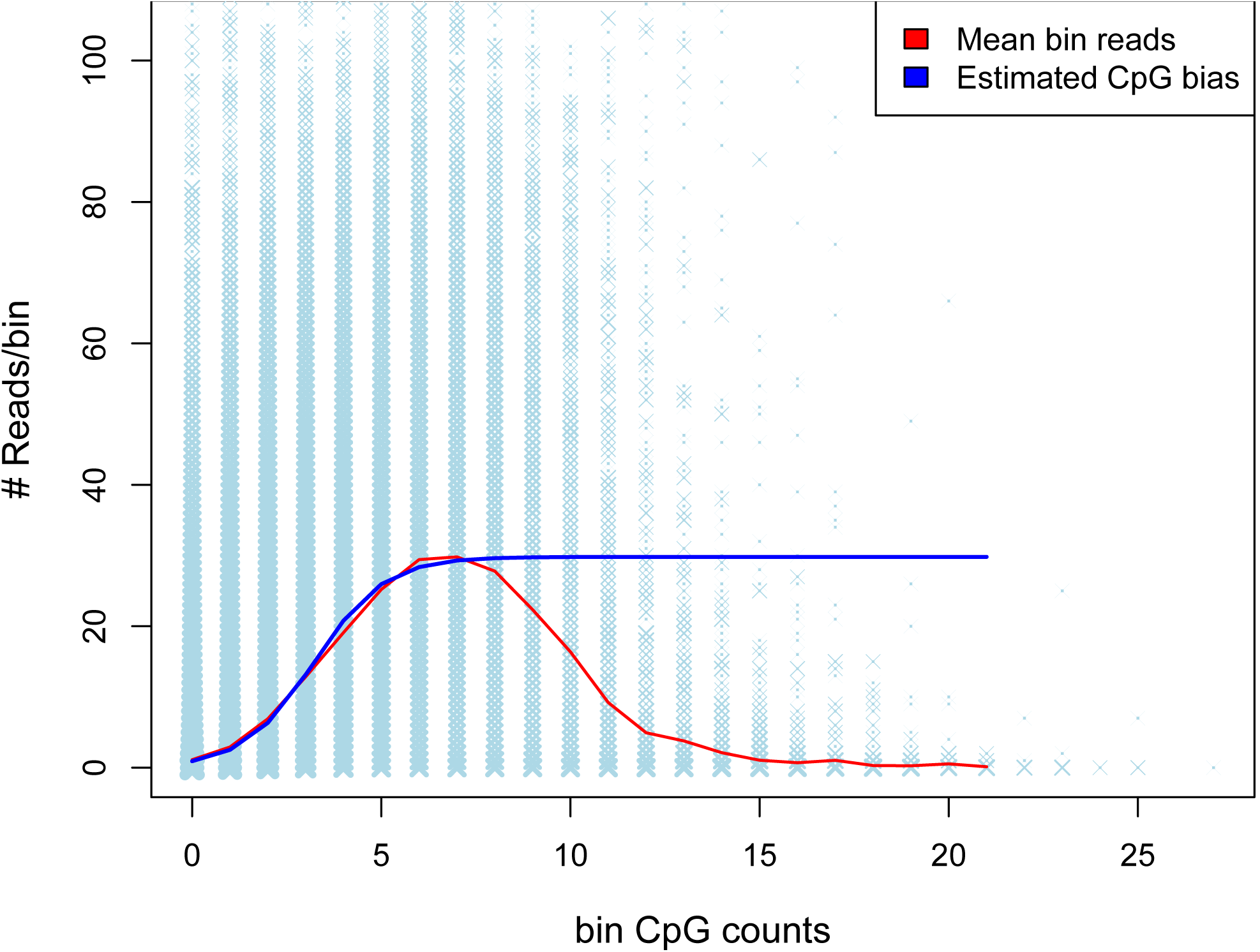
In the ‘MeDEStrand’ method, a logistic regression model was fitted to estimate CpG bias from the raw signals (a.k.a. observed bin reads).

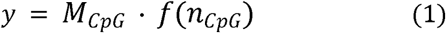

The CpG density effect *f*(*n*_*CpG*_) is to be estimated from the MeDIP-seq signals and eliminated from the observed signals by division from both sides of (1). Then we have

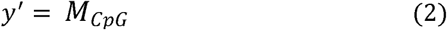

Heuristically, we found log transformation before scaling further improves the accuracy, we log transformed y’ and have y”:

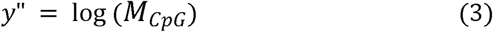

We normalize *y*” by 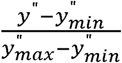 to values between 0 to 1 as the absolute methylation levels for the bins.

Taken into consideration the asymmetric CpG methylation influence on the methyl-CpG DNA enrichment for the positive and negative DNA strands, the estimation and correction of the CpG bias is performed for the positive and negative DNA strands separately. Then, the mean of the inferred absolute methylation levels from both strands is reported as the inferred genome-wide absolute methylation level.

The steps of our method are described as follows:

### The Algorithm workflow

**Input**: MeDIP-seq data

**Output**: bin-based absolute methylation levels

**For** each DNA strand **do**

Divide the given chromosome(s) into user-specified bp bin size (recommend 50 or 100 bp) and count bin reads for the positive DNA strand and the negative DNA strand separately.

1) Re-organize all the bins into groups of same CpG counts and put them in the ascending order according to the bin CpG counts.
2) Let the bin CpG count n_CpG_ of a group be the explanatory variable and the mean bin reads of the group 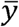 be the response variable. Fit the logistic regression model:

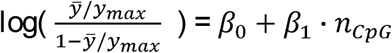 *y*_*max*_: the maximum of 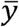
3) Divide the bin reads by the corresponding estimated CpG density effect

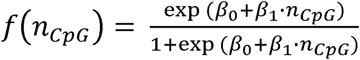

from the fitted model in 2).
4) Log transform the corrected bin reads
5) Scale the bin reads from 4) to values between 0 to 1, and report them as the inferred strand-specific bin-based absolute methylation levels.

Merge the inferred bin absolute methylation values from the positive and negative DNA strands by taking the mean of the corresponding bins. Report them as genome-wide bin-based absolute methylation levels.

**End**

## Results

### Criteria for evaluation

To evaluate the accuracy of the inferred methylation levels, we used the RRBS data as the ‘gold standard’. More specifically, we compared the inferred values with the methylation levels of CpGs fell within the bins from the RRBS data. We calculated the mean of the RRBS CpG methylation in each bin as the ‘true’ methylation levels for the bin/locus. From the ENCODE protocol, each CpG from the RRBS data was covered by at least 10 reads. So, we keep all the RRBS CpGs without further filtering, since 10 reads coverage gives good confidence. For the validation, we kept bins within which there are at least 4 RRBS CpGs, as this provides methylation information from at least two non-adjacent CpGs (Figure 2).

We used Pearson correlation coefficient (PCC) and Spearman correlation coefficient (SCC) as the criteria to measure the agreement between the inferred methylation levels from the MeDIP-seq data and the ‘true’ methylation levels derived from the RRBS data as above. PCC and/or SCC were used as the primary criteria for method evaluation and comparison in the previous studies [2, 5, 7, 9, 10]. While PCC assesses linear relationships between two sets of data, SCC uses the ranks of the values and assesses monotonic relationships (whether linear or not). Higher PCC and/or SCC indicates higher concordance between the inferred absolute methylation levels and the RRBS data. We included the two criteria to make the comparison on a fairer base and the conclusions more reliable.

### Comparison with other methods

To assess the performance of our method ‘MeDEStrand’, we compared it with three other state-of-the-art methods ‘MEDIPS’, ‘BayMeth’ and ‘QSEA’. For each method, we chose the version(s) that could be run on the available data (i.e., MeDIP-seq) to infer absolute methylation levels. For ‘BayMeth’, we used the version ‘SssI-free’, since a fully methylated experimental control sample (e.g., DNA treated with SssI CpG methyl-transferase) preferred by ‘BayMeth’ is not available. For ‘QSEA’, there are three versions. We chose two of them, ‘QSEA(TCGA)’ and ‘QSEA(blind)’. The third version also requires a fully methylated experimental control sample, which is not available. By contrast, ‘QSEA(TCGA)’ and ‘QSEA(blind)’ constructed virtual control samples such that no additional experimental data is required. QSEA(‘TCGA’) constructed its virtual control sample using curated information from the previous 172 samples of the TCGA lung cancer study [21, 22], generalized the results to other cell types. QSEA(‘blind’) constructed its virtual control sample from approximated relationship based on current knowledge. For ‘MEDIPS’ and our method ‘MeDEStrand’, no additional experiment data is required. Some other methods such as ‘methylCRF’ cannot be included, as ‘methylCRF’ was designed for paired-end sequencing reads while the data from ENCODE are those of single-end sequencing reads.

We ran all the methods on the 22 chromosomes (from chromosome 1 to 22) of the MeDIP-seq data to infer absolute methylation levels for the comparison. Among the four cell types, GM12878, K562 and mammary epithelial are cells from female donors and foreskin fibroblast is from male donors. So, we did not include chromosome Y. We did not include chromosome X either since we found chromosome X of GM12878 MeDIP-seq data is corrupted.

‘Bin size’ is an important parameter, which deters the resolution of the inferred absolute methylation levels. In the previous studies, 50 bp or 100 bp bin size were deemed as ‘high resolution’ besides ‘single-base resolution’ results. Since all the methods in the comparison infer bin-based absolute methylation levels, we chose ‘bin size’ of 25bp, 50bp and 100bp to show the performance of the methods.

Figure 5 illustrates the performance of each method measured by PCC between the inferred absolute methylation levels with the RRBS data. Panel A, B on the top and panel C, D at the bottom correspond to the results for two immortalized cell lines GM12878 and K562 and two primary cells foreskin fibroblast and mammary epithelial, respectively.

**Figure 5.**
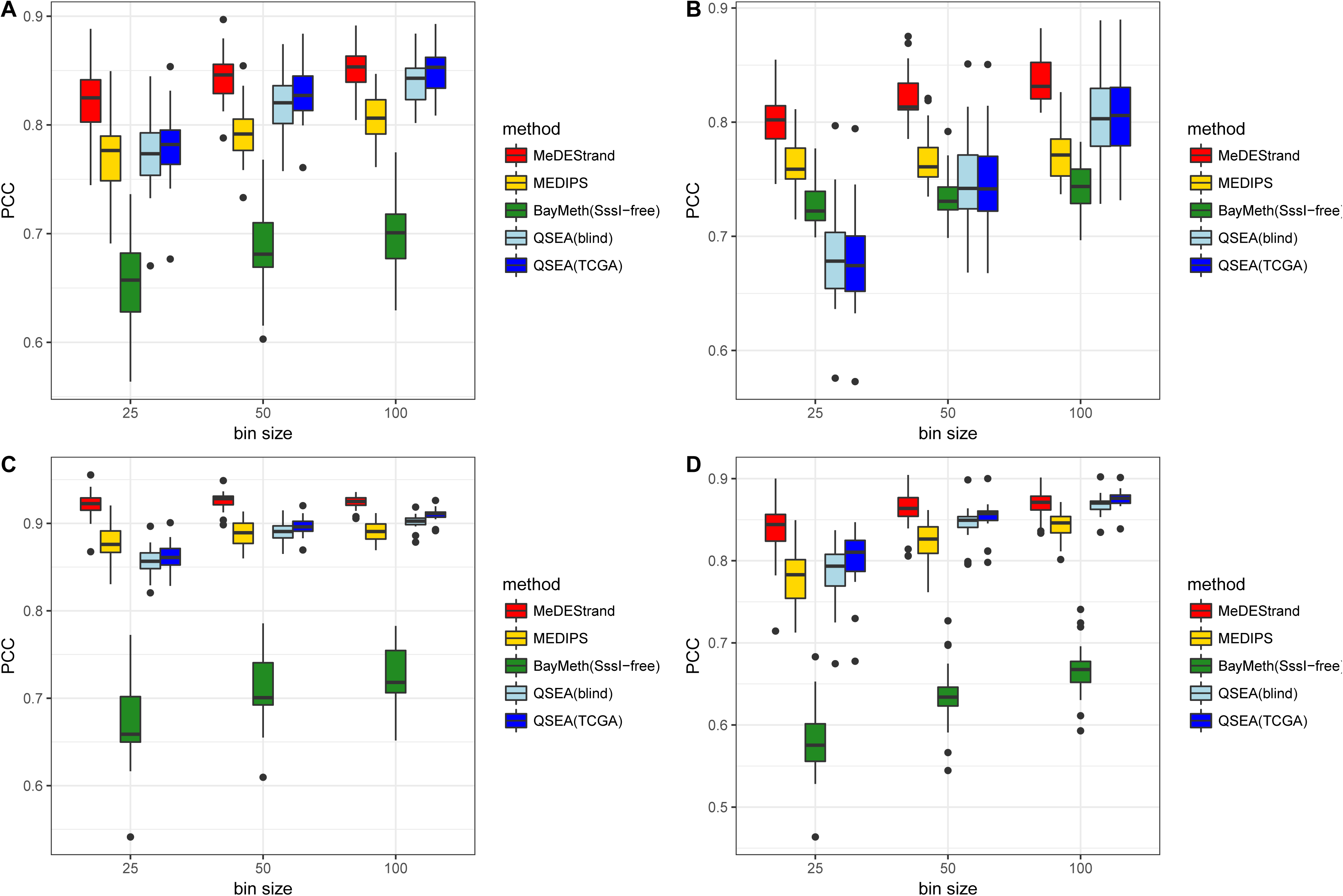
Comparison of the performances of four methods by Pearson correlation coefficient on four cells. A: GM12878. B: K562. C: foreskin fibroblast. D: mammary epithelial. Y-axis shows the values of Pearson correlation coefficient. X-axis shows the parameter ‘bin size’ varying from 25bp to 100bp. Boxplot illustrates the variation of Pearson correlation coefficients across the 22 chromosomes.

It is observed that ‘BayMeth’ does not perform well at any bin size for all the cell types selected. This may be due to the lack of a full-methylated experimental control sample which is preferred by the ‘BayMeth’ model to make good inference. Considering the results from all the cell types and bin sizes, it can be concluded that ‘MeDEStrand’ has the best performance with regards to the median value of PCCs across the 22 chromosomes. We also notice ‘QSEA(blind)’ and ‘QSEA(TCGA)’ have comparable performance with ‘MeDEStrand’ at bin size 100bp, however, with a larger variation of PCCs in general. For cell type mammary epithelial, ‘QSEA(TCGA)’ has slightly better performance than ‘MeDEStrand’.

Figure 6 has the same layout as Figure 5 and illustrates the performance of each method measured by SCC. It can be noticed that according to the SCC criterion, ‘BayMeth’ has significantly improved performance while ‘QSEA(blind)’ and ‘QSEA(TCGA)’ have reduced performance at bin size 25bp and 50bp. Meanwhile, ‘MeDEStrand’ remains as the best performer among all the methods at all bin sizes for all the cell types.

**Figure 6.**
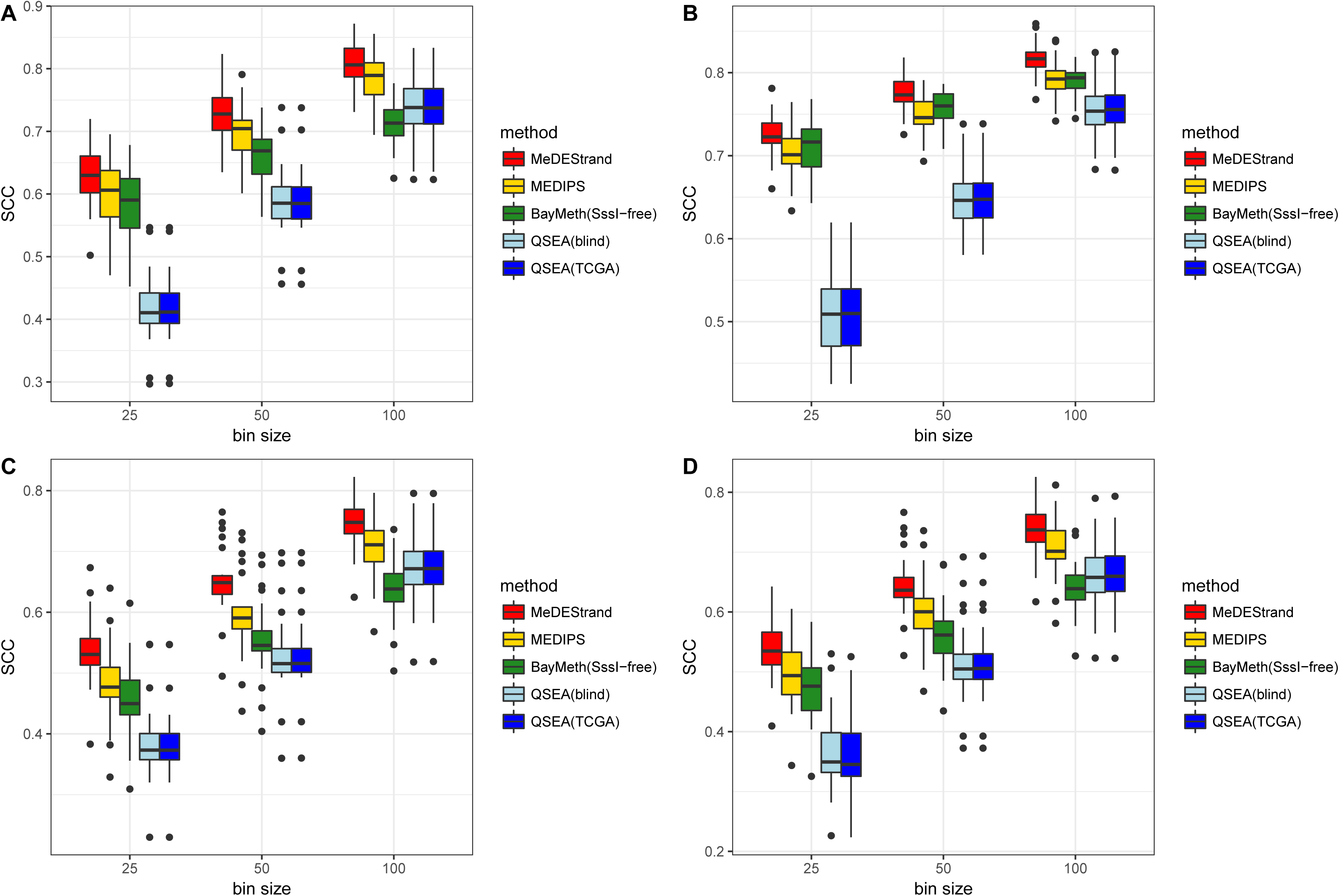
Comparison of the performances of four methods by Spearman correlation coefficient on four cells. A: GM12878. B: K562. C: foreskin fibroblast. D: mammary epithelial. Y-axis shows the values of Spearman correlation coefficient. X-axis shows the parameter ‘bin size’ varying from 25bp to 100bp. Boxplot illustrates the variation of Spearman correlation coefficients across the 22 chromosomes.

Moreover, from Figure 5 and Figure 6, we can see in general, ‘MeDEStrand’ not only provides the highest median values of PCCs and SCCs for the 22 chromosomes evaluated, but also, has smaller variation in PCCs and SCCs across the 22 chromosomes. This result shows the stable performance of ‘MeDEStrand’ across the 22 chromosomes.

Finally, the processing time for the whole genome (including the time to import the data) ranges from ∼ 15 minutes to ∼ 3 hours based on a MacBook Pro laptop with 16G RAM depending on the method. ‘MeDEStrand’ has one of the shortest processing time among all the methods (i.e., ∼15 minutes). We note that the processing time is not a key criterion for method comparison since all the methods provide reasonably fast processing.

Taken together, we have demonstrated that ‘MeDEStrand’ is one of the best and robust methods to infer genome-wide absolute methylation levels at bin size 25bp, 50bp and 100bp. Smaller bin size corresponds to higher resolution, which is desirable. In addition to the inference of genome-wide absolute methylation levels, ‘MeDEStrand’ also provides differential methylation analysis based on the inferred absolute methylation levels between two sample groups. ‘MeDEStrand’ has been implemented as a R package and is freely available for download from GitHub: https://github.com/jxu1234/MeDEStrand.git

### Improvement from using a sigmoid function to estimate CpG bias and the strand-specific reads processing

As described previously, ‘MeDEStrand’ uses a sigmoid function to estimate CpG bias from the DNA enrichment signals. In addition, ‘MeDEStrand’ estimates and corrects CpG bias from the enrichment signals for the positive and negative DNA strands separately, and reports the average of the inferred strand-specific absolute methylation levels as the genome-wide absolute methylation levels. We are interested to know if these two modifications contribute to the overall improvement, respectively.

To answer the question, we constructed a modified version, namely, ‘MEDIPS(strand-processing)’, which has the same algorithm as ‘MEDIPS’ except added the procedure of processing reads mapped to the positive and negative DNA strands separately. To illustrate the gain at different CpG density regions, we divided all bins into four categories based on their CpG counts. The first category consists of bins with CpG counts from the minimum to the 1^st^ quartile and corresponds to the ‘low’ CpG density regions. The second category consists of bins with CpG counts from the 1^st^ quartile to the median and corresponds to the ‘lower-medium’ CpG density regions. The third category consists of bins with CpG counts from the median to the 3^rd^ quartile and corresponds to the ‘higher-medium’ CpG density regions. The last category consists of bins with CpG counts from the 3^rd^ quartile to the maximum and corresponds to the ‘high’ CpG density regions. These four categories represent different DNA CpG density patterns within the bins. We calculated PCCs for the categories using cell line GM12878 at bin size 100 bp for the illustration.

Figure 7 shows the performance of ‘MEDIPS’, ‘MEDIPS(strand-processing)’ and ‘MeDEStrand’ at different CpG density regions. We observe that ‘MEDIPS(strand-processing)’ had improved performance compared to ‘MEDIPS’ at all CpG density regions. The result shows that merely adding the procedure to process reads mapped to the positive and negative DNA strands separately improves the overall performance of ‘MEDIPS’. We note that for all the previous methods, bin reads were counted by combining reads mapped to the same loci regardless of their strand information. However, we demonstrated that the strand-specific information can be utilized to improve the performance. Although depending on the models, assumptions and techniques to make inference, it may not necessarily be utilized in the same way as we did.

**Figure 7.**
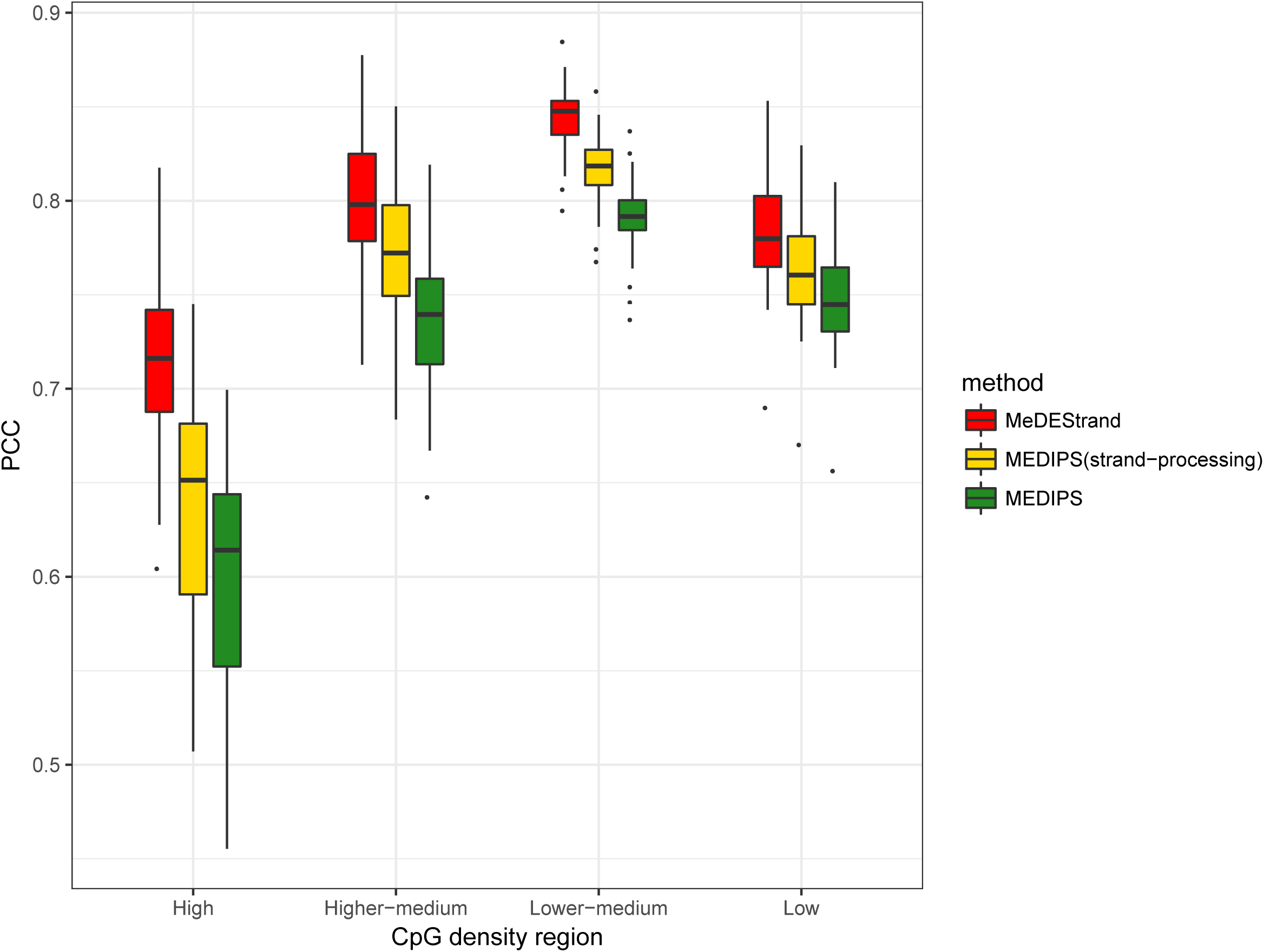
Comparison of ‘MeDEStrand’ with ‘MEDIPS’ and its modified version ‘MEDIPS(strand-processing) at different CpG density regions. The MeDIP-seq and RRBS data of GM12878 were used for the illustration.

Our analysis also revealed that the CpG density region ‘lower-medium’ has the highest PCCs for all the methods and the PCCs decrease from this category when the regional CpG density decreases or increases. This may be because at ‘low’ CpG density regions, the inference of the absolute methylation levels is often more difficult for the enrichment-based methods than for the bisulfite conversion methods. At lower sequencing depth, the lack of methylation cannot be distinguished from the lack of coverage. This is due to the stochastic nature of read coverage from the enrichment-based methods. At the ‘high’ CpG density regions, ‘MEDIPS’ and ‘MEDIPS(strand-processing)’ had significantly deteriorated performance with their median PCC reduced to ∼ 0.6. We also see the largest variation of PCCs across the 22 chromosomes for those two methods. By comparison, ‘MeDEStrand’ has the median of PCCs above 0.7 with much smaller variation. This category corresponds to the high CpG density regions and it is the category where ‘MeDEStrand’ made the most improvement compared to ‘MEDIPS’. The improvement demonstrates the advantage of a sigmoidal logistic regression model over a linear regression model to estimate the CpG bias, with its upper asymptote accommodating for the saturation effect of methyl-CpG binding. Still, it is noted at the high CpG density regions, all the methods make less accurate inference compared to the results at the other CpG density regions.

Figure 7 also illustrates the synergistic effects for improvement by utilizing the strand-specific information combining with the sigmoid function to estimate and correct CpG bias, i.e., ‘MeDEStrand’ showed better performance than ‘MEDIPS(strand-processing)’ at all CpG density regions.

#### Improvement from strand-specific information

To inspect how strand-specific information improves the overall performance, we compared ‘MEDIPS’ and ‘MEDIPS(strand-processing)’. The latter differs from the former only by the additional procedure to process reads mapped to the positive and negative DNA strands separately. ‘MEDIPS’ counts bin reads by combining reads mapped to the same genomic loci regardless of their strand information, i.e., the bin reads for the loci is the total number of reads fell in the bin from both DNA strands. For ‘MEDIPS(strand-processing)’, reads mapped to the positive and negative DNA strands are counted by the bins for the positive and negative DNA strands separately, i.e., strand-specific bin reads. The corresponding strand-specific bins at the same loci may or may not have the same bin reads (Figure 2). We observed ∼ 40% of genomic coverage has different strand-specific bin reads. We wonder whether it is the asymmetric strand-specific bin reads or it is merely the procedure to estimate and correct CpG bias for the positive and negative DNA strands separately (i.e. irrelevant to the asymmetry of the bin reads) that contributes to the improvement.

To investigate the question, we devised a counting scheme that eliminates the asymmetry of the bin reads from ‘MEDIPS(strand-processing)’. That is, we divide each bin reads from ‘MEDIPS’ evenly and re-assign the halved bin reads to the corresponding bins residing on the positive and negative DNA strands as the bin counts for ‘MEDIPS(strand-processing)’. Note this re-assignment has no effects on the results of ‘MEDIPS’, since combined reads from both DNA strands remain the same. However, for ‘MEDIPS(strand-processing)’, the asymmetry of the bin reads got lost. After we re-ran ‘MEDIPS(strand-processing)’, we observed that there is no improvement from ‘MEDIPS(strand-processing)’; even ‘MEDIPS(strand-processing)’ still incorporates the procedure to estimate and correct CpG bias for the positive and negative DNA strands separately. In fact, ‘MEDIPS’ and ‘MEDIPS(strand-processing)’ methods now have the same performance. The results show that ‘MEDIPS’ can be viewed as a special case of ‘MEDIPS(strand-processing)’, where strand-specific bin reads for the positive and negative DNA strands are treated as equal. By combing the reads from both DNA strands as the bin reads for the loci, the strand-specific information was discarded automatically.

However, we demonstrated the improvement from the strand-specific processing based on the strand-specific bin reads counted for the positive and negative DNA strands separately (Figure 7, ‘MEDIPS(strand-processing)’ vs. ‘MEDIPS’). The asymmetric bin reads preserve certain strand-specific methylation information that can be used to improve the overall performance of the methods.

## Discussions

CpG bias is a major confounding factor that affects the inference of absolute methylation level from the enrichment-based DNA methylation profiling data. In this work, we developed a method that utilizes a logistic regression model to fit the sigmoidal increment of Methyl-CpG binding signal associated with the CpG density. The estimated relationship is used as CpG bias in our model to correct the raw signals (a.k.a. observed bin reads). Our method ‘MeDEStrand’ was evaluated based on the MeDIP-seq and RRBS data of two immortalized cell lines GM12878 and K562, and two primary cells foreskin fibroblast and mammary epithelial. The comparison study demonstrated the improved performance of our method under both criteria of PCC and SCC. Besides MeDIP-seq data, ‘MeDEStrand’ can also be applied to other enrichment-based sequencing data generated using MethylCap-seq/MDB-seq in which the main bias also comes from CpG density. However, we are unable to evaluate ‘MeDEStrand’ on these types of data since matching pairs of RRBS and MethylCap-seq/MDB-seq are not available in the public repository.

We showed that separate processing of the sequencing reads mapped to the positive and negative DNA strands could improve the performance of the methods. In the antibody capture method (e.g., MeDIP-seq), the genomic DNA is denatured and fragmented before the single-stranded methylated DNA fragment is pulled down by immunoprecipitation procedure [1, 23, 24]. In this way, the fragments carry strand information. Whereas in the Methyl-CpG-binding domain (MBD) protein capture methods, such as MethylCap-seq/MBD-seq, the genomic DNA is not denatured, and the double-stranded methylated DNA fragment is captured before sequencing therefore technically they do not carry strand information [24, 25]. We are unclear whether the procedure to process reads in the strand-specific way can also improve the performance for MethylCap-seq/MBD-seq data.

Further investigation can be conducted if matching pairs of MethylCap-seq/MBD-seq and RRBS data are available.

Although ‘MeDEStrand’ showed improved performance compared to ‘MEDIPS’ at all the CpG density regions, less accurate results were observed at high CpG density regions (Figure 7). Future work will identify the cause and improve the inference of absolute methylation level for these regions.

Previous methods showed performance gain from explicitly modeling of copy number variation (CNV), which directly affects read density [24, 26]. In a 2013 paper, 37 available tools were reviewed to identify whole-genome CNVs based on various computational strategies [27]. Further improvement may be possible by incorporating a suitable one to our current work.

DNA methylation occurs at C5 position of cytosine within CpG dinucleotides and non-CpG cytosine in plants and embryonic stem cells in mammals [28, 29]. For the somatic cell lines, DNA methylation dominantly occurs at the CpG sites. Whereas for the embryonic stem cells, 25% of the DNA methylation occur at the CHG and CHH sites [30]. Unlike the enrichment methods based on Methyl-CpG-binding domain (MBD) protein, which only binds to the double-stranded DNA methylated CpG sites, antibody-based MeDIP-seq method also captures CHG and CHH methylation sites. Current methods that infer DNA absolute methylation only considered the CpG methylation effects for the enrichment [6-11]. To our best knowledge, there is no method that has incorporated CHG and CHH methylation effects in the model. For embryonic stem cells or those cells where a significant amount of DNA methylation occurs at non-CpG sites, CHG and CHH methylation may be taken into consideration for further improvement in the inference of DNA absolute methylation levels from MeDIP-seq data.

## Conclusions

‘MeDEStrand’ outperforms other state-of-the-art methods at high resolutions (25bp-, 50bp- and 100 bp-sized bins) based on the evaluations on the four independent datasets.

In addition, ‘MeDEStrand’ reaches a comparable level of accuracy relying on MeDIP-seq data only. In contrast, ‘BayMeth’ and ‘methylCRF’ require additional experiment data to achieve high accuracy. Thus, ‘MeDEStrand’ is a useful tool to analyze data in the public repository where additional experiment data are provided.

Finally, estimating and correcting CpG bias by a sigmoidal model as well as processing reads in the strand-specific way was shown to improve the overall performance. This direction of exploring strand-specific information for improvement can be considered in future method development.

## Abbreviations

WGBS: whole-genome bisulfite sequencing
RRBS: reduced-representation bisulfite sequencing
PCC: Pearson correlation coefficient
SCC: Spearman correlation coefficient
bp: base pair
kb: A kilobase pair, i.e., 1000 base pairs

## Ethics approval and consent to participate

Not applicable

## Consent for publication

Not applicable

## Availability of data and material

The datasets analyzed during the current study are publicly available from ‘Experiment Matrix-ENCODE’ repository from the link (https://www.encodeproject.org/matrix/?type=Experiment&status=released&assay_slims=DNA+methylation&replicates.library.biosample.donor.organism.scientific_name=Homo+sapiens%29&replicates.library.biosample.donor.organism.scientific_name=Homo+sapiens)

## Authors’ contributions

JX and YD conceptualized the model. JX implemented the model and wrote the R package. JX and YD conducted model comparison and drafted the manuscript. SW, PY and SB contributed result evaluation and manuscript revising. All authors have approved the final manuscript.

## Acknowledgements

Not applicable

## Author’s information

^1^ Department of Bioengineering, University of Illinois at Chicago, Chicago, Illinois, United States of America. ^2^ Division of Reproductive Science in Medicine, Department of Obstetrics and Gynecology, Feinberg School of Medicine, Northwestern University, Chicago, Illinois, United States of America

Jingting Xu^1^: Yang Dai^1^

Shimeng Liu^2^: Ping Yin^2^

Serdar Bulun^2^

